# Antigen-Specific T Cell Recall Assay To Screen Drugs For Off-Target Effects

**DOI:** 10.1101/2020.08.10.241869

**Authors:** Eden Kleiman, Gloria Sierra, Dennie Magcase, Marybeth V George, Pirouz M Daftarian

## Abstract

Extracellular adenosine suppresses T cell immunity in the tumor microenvironment. We have developed an in vitro recall assay utilizing a sequential adenosine dosing regimen guide for testing drug off-target effects on memory T cell expansion/function. As a proof of principle, we show low dose adenosine analog GS-5734, a monophosphoramidate prodrug of an adenosine analog, does not alter memory T cell recall whereas toxicity observed at high dose favors antigen-specific memory T cell survival/proliferation over non-specific CD8^+^ T cells. Parent drug GS-445124 at high dosage interferes with antigen-specific T cell recall without cellular toxicity. Despite similar chemical structure, these drugs displayed opposing effects on memory T cell expansion. This assay platform has broad utility in screening memory T cell off-target effects.

## INTRODUCTION

The recent emergence of SARS-CoV-2 has redirected research efforts towards understanding the adaptive immune response and enhancing vaccine development targeting this virus. Recent evidence reinforces the crucial role of a strong T cell immune response in SARS-CoV-2 disease ^1,2^ control while also illustrating the link between disease progression and the inability to sequester appropriate T cell help ^3,4^. GS-5734 (Remdesivir) is a monophosphoramidate prodrug of an adenosine analog currently being utilized for therapeutic use in SARS-CoV-2 infections ^5^ due to its ability to potently inhibit SARS-CoV-2 viral load by blocking viral activity of RNA-dependent RNA polymerase (EC_50_ range of 0.77μM to 23.15μM in SARS-CoV-2 infected Vero E6 cells) ^6–8^. Remdesivir has previously shown effectiveness against MERS, SARS, Ebola and other viruses ^9^ and can prevent SARS-CoV-2-mediated progression to pneumonia in rhesus macaques ^10^. It is unclear if the antiviral effects of adenosine-related drugs are related to any potential modulation of T cell immunity as a mechanism of action or off-target effect.

Interestingly, adenosine itself is a suppressive agent in the tumor microenvironment (TME) that depresses cytotoxic T cell activity. Cancer-related hypoxia can elevate extracellular adenosine due to enzymatic ATP catabolism, particularly by the rate limiting ectonucleotidase enzyme CD73. Local fluctuations in adenosine can cause TME concentrations to reach into the hundreds of micromolars ^11, 12^. Extracellular adenosine exerts its immunosuppressive effects primarily as a ligand for A2A adenosine receptor (A2AR) which upregulates intracellular cAMP. Genetic deletion or pharmacologic antagonism of A2AR enables CD8^+^ T cell-dependent tumor rejection but can also exacerbate inflammatory induced tissue damage ^12^. In addition to the TME, adenosine has been reported to be involved in a myriad of other pathologies such as Parkinson’s disease, fibrosis, hepatic steatosis, colitis, asthma and diabetes ^13^. Given the importance of T cell mediated immunity in counteracting viral infections such as SARS-CoV-2, assays to screen small molecule drugs (e.g. anti-virals) for potential off-target effects on memory T cells would be helpful. We have developed an in vitro assay that uses sequential adenosine additions to suppress memory T cell expansion as a means to test adenosinergic pathway inhibitors (unpublished internal data). Here we present a repurposed version of this assay using drugs similar in structure to adenosine to test off-target drug effects on antigen-specific T cell recall and functionality. Remdesivir at low doses does not inhibit antigen-specific recall but toxicity is observed at high doses consistent with the reported CC50 ^7^, and coinciding with favored expansion/survival of antigen-specific T cells over the general CD8^+^ T cell pool. Surprisingly, parent drug GS-441524 displayed inhibitory effects on antigen-specific T cell recall at high dosage with no adverse effect on general cell viability. Using surface CD137 expression as an activation read-out, high dose Remdesivir increases antigen-specific T cell activation while parent drug caused elevated expression in EBV peptide treated samples. Thus, this in vitro assay can be used broadly as a quick preliminary screen for anti-viral drugs and other small molecule drugs with respect to their potential effect on both CD8^+^ as well as CD4^+^ T cell memory response.

## MATERIALS AND METHODS

### Recall assay

Donor PBMCs (Astarte Biologics, now part of Cellero, Lowell, MA) were plated at 500,000 cells/100uL complete media per well in a round-bottom 96-well plate. Complete media comprised AIM V medium (ThermoFisher Scientific, Waltham, MA, Cat # 12055091) with 5% human AB serum (Millipore Sigma, Burlington, MA, Cat # H3667). Peptide stimulation in appropriate wells was given at 10μg/mL final concentration unless otherwise stated. NLVPMVATV peptide was purchased from Astarte Biologics. All other peptides were purchased from MBLI (Chicago, IL). For 7-day culture experiments, 100uL per well of culture medium was replaced with 100uL fresh complete media supplemented with 50U/mL IL-2 (Millipore Sigma, Cat # 11147528001) on days 1,3 and 5 unless otherwise stated. Adenosine (Millipore Sigma, Cat # A9251) was given at 1mM final day 1 and 0.5-0.75mM final days 3 and 5 unless otherwise stated. Adenosine was dissolved in complete media unless otherwise stated. For 48-hour stimulation of previously expanded PBMCs, the ratio of expanded to unexpanded is 2:1 with 500,000 cells total per well. GS-5734 and GS-445124 (Cayman Chemical, Ann Arbor, MI) were dissolved in Hybri-Max DMSO (Millipore Sigma, Cat # D2650). Adenosine deaminase (ADA) (Cat #10102105001) and Inosine (Cat #I4125) were both purchased from Millipore Sigma. Inosine was prepared in complete media.

### Bead-based Proliferation Assay

Donor PBMCs were incubated with 5μM CellTrace Blue Cell Proliferation kit (ThermoFisher, Cat # C34574) proliferation dye and then stimulated with anti-CD3/anti-CD28 beads (ThermoFisher, Cat # 11132D) in a 96-well plate. Adenosine was added to appropriate wells Day 0 at 1mM final concentration and again on Day 1 at 0.5mM final concentration. Proliferation was assessed 4 days post-stimulation by flow.

### Surface and intracellular staining

PBMCs were incubated with 50nM Dasatinib (Selleck Chemical LLC, Cat # S1021) 15 minutes prior to antibody/tetramer staining for 30 minutes at room temperature in FACS buffer (0.5% BSA in PBS). Cells were then washed and resuspended in FACS buffer with 7-AAD (MBLI, Code # FP00020050). For Annexin V (BioVision Inc., Milpitas, CA, Cat # K101-100) and 7-AAD co-staining, both were added at the same time in 1X Annexin V binding buffer.

For intracellular stains, cells were washed twice with PBS and then fixed, permeabilized and stained in permeabilization buffer according to manufacturer’s guidelines. Intracellular staining was either performed using the True-Nuclear Transcription Factor Buffer Set (Biolegend, San Diego, CA, Cat # 424401) or CytoFix Fixation buffer / FACS Permeabilizing solution 2 (BD Biosciences, San Jose, CA, Cat # 554655 / 340973). Cells were then washed twice in PBS and resuspended in PBS. For cytokine staining, 5ug/mL brefeldin A (Millipore Sigma, Cat # B7651) was added 4 hours prior to surface staining.

Detection reagents used are as follows: CD3 FITC (Code # FP10255010), CD4 APC (Code # FP10328010), HLA-A*02:01 CMV pp65 tetramer NLVPMVATV-PE (Code # TB-0010-1), T-Select HLA-A*11:04 EBV EBNA3B 416-424 tetramer IVTDFSVIK-PE (Code # TS-M029-1), HLA-A*11:01 EBV EBNA3B tetramer AVFDRKSDAK-PE (Code # TS-M028-1), HLA-A*11:01 CMV pp65 tetramer ATVQGQNLK-PE (Code # TB-M012-1) were from MBLI. CD8 BV510 (Cat # 344732), TIM-3 BV711 (Cat # 345024), CD137 BV421 (Cat # 309820), CD45RO BV785 (Cat # 304234), CD14 BV785 (Cat # 301840), CD11b BV711 (Cat # 301344) were from Biolegend. PD-1 BV421 (Cat # 564323) and IFNγ BV421 (Cat # 562988) were from BD Biosciences. LAG-3 APC (Cat # 17-2239-42), PD-L1 PE (Cat # 12-5983-42) and PD-L1 PE (Cat # 12-5888-42) were from (ThermoFisher - eBioscience). TOX APC (Cat # 130-118-474) was from Miltenyi Biotec (Bergisch Gladbach, Germany).

### Analysis software

Statistical analysis was performed using GraphPad Prism software (GraphPad, San Diego, CA). All bar graph data presented is shown as mean with standard deviation. Flow cytometry analysis was performed using FlowJo software (BD Biosciences).

## RESULTS

We employed a reductionist approach to simulate one aspect of the TME environment by incorporating a series of high dose adenosine treatments during the course of the recall assay to specifically suppress antigen-specific memory T cell expansion without loss of PBMC viability. We adopted a flow cytometry-based tetramer readout as a measure of CD8^+^ antigen-specific T cell expansion on day 7 post-peptide stimulation. Adenosine-mediated suppression of memory T cell expansion (a.k.a. recall) was achieved by adding adenosine on days 1, 3 and 5 (Fig. 1A). This regimen gave the most consistent suppression, although adenosine treatment on days 3 and 5 also produced a robust suppression. We chose to stick with treatment on days 1, 3 and 5 going forward (Fig. 1B). This assay has utility in screening drugs targeting the adenosinergic signaling pathway, particularly A2AR receptor antagonists (unpublished internal data). We have repurposed the assay to screen drugs for off-target memory T cell effects which involve the same dosing regimen as adenosine (Fig. 1B).

**Figure 1.**
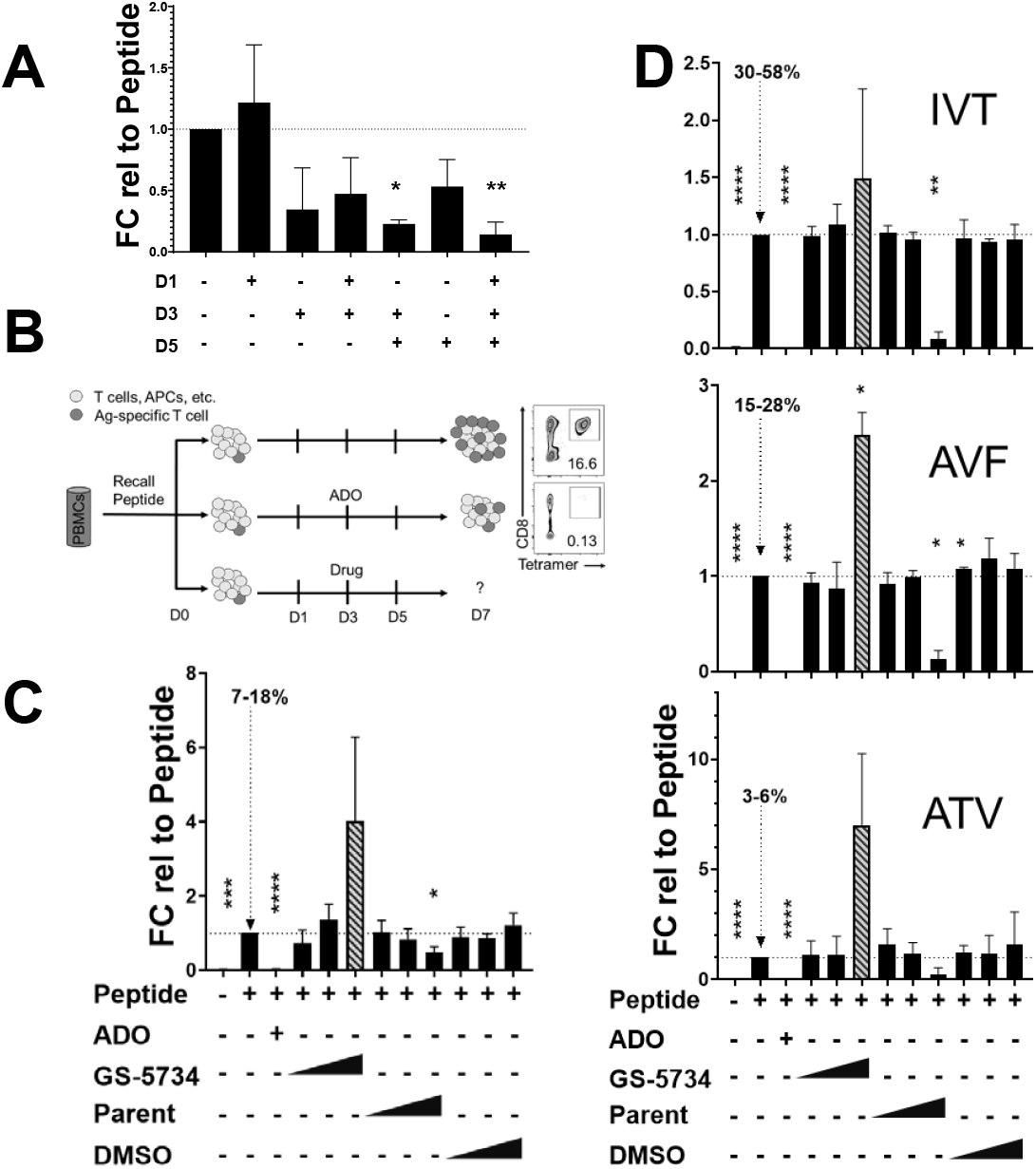
Adenosine-based recall assay. A) HLA-A*02:01 Donor #1 PBMCs treated either with peptide alone (CMV pp65 NLVPMVATV) or peptide plus adenosine (ADO) on indicated days and assayed on day 7. All initial ADO treatments were 1mM final concentration while all secondary or third treatments are at a final ADO concentration of 0.5-0.75mM. Data are representative of 3 independent experiments, except for the D3 (n of 2) and displayed as percent tetramer fold-change (FC) relative to peptide-only treated cells (comparison using horizontal dotted line) within each respective experiment. B) Assay setup whereby donor PBMCs are treated with ADO days 1, 3 and 5 and assayed for CD8+ tetramer positive cells. ADO results in significantly reduced tetramer positive T cell expansion. Drug arm uses the same dosing regimen as ADO. C) Donor #1 PBMCs were treated with NLVPMVATV or D) HLA-A*11:01 Donor #3 PBMCs treated with EBV EBNA-3B peptide IVTDFSVIK, EBV EBNA4 peptide AVFDRKSDAK, or CMV pp65 peptide ATVQGQNLK. The percent positive range of tetramer positive cells (among live T cells) for peptide-only stimulated wells is indicated with dotted arrow. GS-5734 and GS-445124 were used at increasing doses of 500nM, 5μM and 50μM. 50μM GS-5734 treatment is indicated with striped bar to denote its high toxicity. DMSO was administered at the same concentrations as drugs; 0.1%, 0.01% or 0.001% final. Data are representative of 5 (C) or 3 (D) independent experiments. Dunnett’s multiple comparison test. *, p ≤ 0.05; **, p ≤ 0.01; ***, p ≤ 0.001; ****, p ≤ 0.0001.

Initially, we sought to characterize the mechanism of adenosine-mediated suppression. Adenosine inhibits proliferation of activated cells as evidenced by suppression of anti-CD3/anti-CD28 activated cells (Fig. S1) as well as suppression of the more physiological stimulation, recall peptide (Fig. 1 and below). Antigen-specific T cells that did emerge from adenosine on day 7 stained positive to the same degree for proliferation marker Ki67 (Fig. S2) indicating that they are capable of proliferating but are less likely to. Further, adenosine in our system did not adversely affect cell viability (below). This indicates adenosine treatment reduces the proliferative capacity of PBMCs in culture. Adenosine metabolites are unlikely to be mediating the suppression as enzyme supplementation with adenosine deaminase (ADA) restored recall and treatment with adenosine breakdown metabolite inosine (at equivalent concentrations) did not show any suppressive effect and in fact slightly increased recall (Fig. S3) consistent with recent evidence that CD8^+^ T cells can use inosine as an alternative carbon source to fuel expansion ^14^. Co-inhibitory receptor surface expression of PD-1 and LAG-3 is not significantly altered on day 7 tetramer positive T cells due to adenosine, whereas TIM-3 is downregulated. Bystander CD8^+^ T cell (tetramer negative) activation in all three markers is lower as a result of adenosine (Fig. S4). Furthermore, short-term treatment with just adenosine (no peptide) significantly increased the percentage and surface expression of PD-1 ligand PD-L1/PD-L2 in CD14^+^ CD11b^-^ cells (Fig. S5). This suggests that T cell activation could be sub-optimal in the presence of adenosine. However, adenosine did not negatively affect short term IFNγ cytokine production or induction of polyfunctional human effector memory CD8^+^ T cell marker TOX (Fig. S6)^15^ on tetramer positive T cells. Thus, whether adenosine actually suppresses T cell activation or suppresses T cell activation enough to affect proliferation requires further investigation. Thus, the major effect of adenosine in this assay system is suppression of proliferation with an as yet undefined mechanism of action.

Recent evidence has suggested GS-5734 as a promising drug in the fight against SARS-CoV-2. We chose to test this prodrug and its parent drug since they were similar in structure to adenosine. We applied the same dosing regimen that was used in sequential adenosine additions wells to assess any potential adverse effects on memory T cell expansion. Data from HLA-A*02:01 donor #1 treated PBMCs with CMV pp65 NLVPMVATV peptide shows that 500nM and 5μM GS-5734 did not alter antigen-specific T cell expansion (Fig. 1C and S7). GS-5734 treatment at 50μM was associated with toxicity, consistent with its reported CC50 ^7^, particularly towards CD8^+^ T cells (Fig. S8-9). However, 50μM GS-5734 resulted in increased antigen-specific T cell percentage relative to non-antigen-specific CD8^+^ T cells. Whether this is due to an increased survival advantage of antigen-specific T cells or increased proliferation in the absence of competing CD8^+^ T cells is unclear. Parent drug GS-445124 did not alter T cell proliferation at 500nM and 5uM but did significantly reduce expansion at 50μM (Fig. 1C) without cellular toxicity (Fig. S9). Donor #1 PBMCs were also treated with nonspecific adenosine receptor agonist NECA and the A2AR-specific agonist CGS-21680. Both had negligible effects on proliferation (Fig. S7) at the doses used. The results using NECA and CGS-21680 are similar to a study reporting marginal effects on murine T cell proliferation ^16^.

We tested another donor, HLA-A*11:01 donor #3, to see the if the effect of these drugs would be consistent, including different antigen peptides. Three peptides were used in separate stimulations; EBV EBNA-3B peptide IVTDFSVIK, EBV EBNA4 peptide AVFDRKSDAK and CMV pp65 peptide ATVQCQNLK. In all cases GS-5734 at high dosage caused significant toxicity, yet consistently increased the percentage of antigen-specific T cells among the T cell pool (Fig. 1D). Also consistent with the previous donor, high dosage of GS-445124 lowered antigen-specific T cell recall (Fig. 1D) without affecting cell viability (Fig. S10).

In addition to memory T cell expansion, off-target effects on antigen-specific memory T cell activation can be assayed using this platform. Day 7 expanded antigen-specific T cells were assessed for surface expression of CD137 which is an inducible co-stimulatory molecule. We chose to focus on CD137 due to its ability to serve as a general marker for antigen-specific T cell activation regardless of cytokine secretion profile and T cell differentiation status ^17^ (Fig. S11). The effect of drug on activation-induced CD137 expression varied by recall antigen used. 50μM GS-5734 increased CD137 expression with all recall peptide stimulations, whereas 50μM GS-441524 combined with EBV peptide stimulations (Donor #3) led to increased CD137 (Fig 2 and S12). CD137 induction in bystander non-antigen specific CD8^+^ T cells was varied. 50μM GS-5734 increased CD137 surface expression in half of the cases but decreased CD137 expression in ATVQGQNLK treated cells, whereas the effect with 50μM GS-441524 was mostly decreased CD137 expression (Fig. 2). Treatment of ATV peptide-stimulated cells highlights the disparate effects that a drug can have on antigen-specific T cells versus bystander T cells.

**Figure 2.**
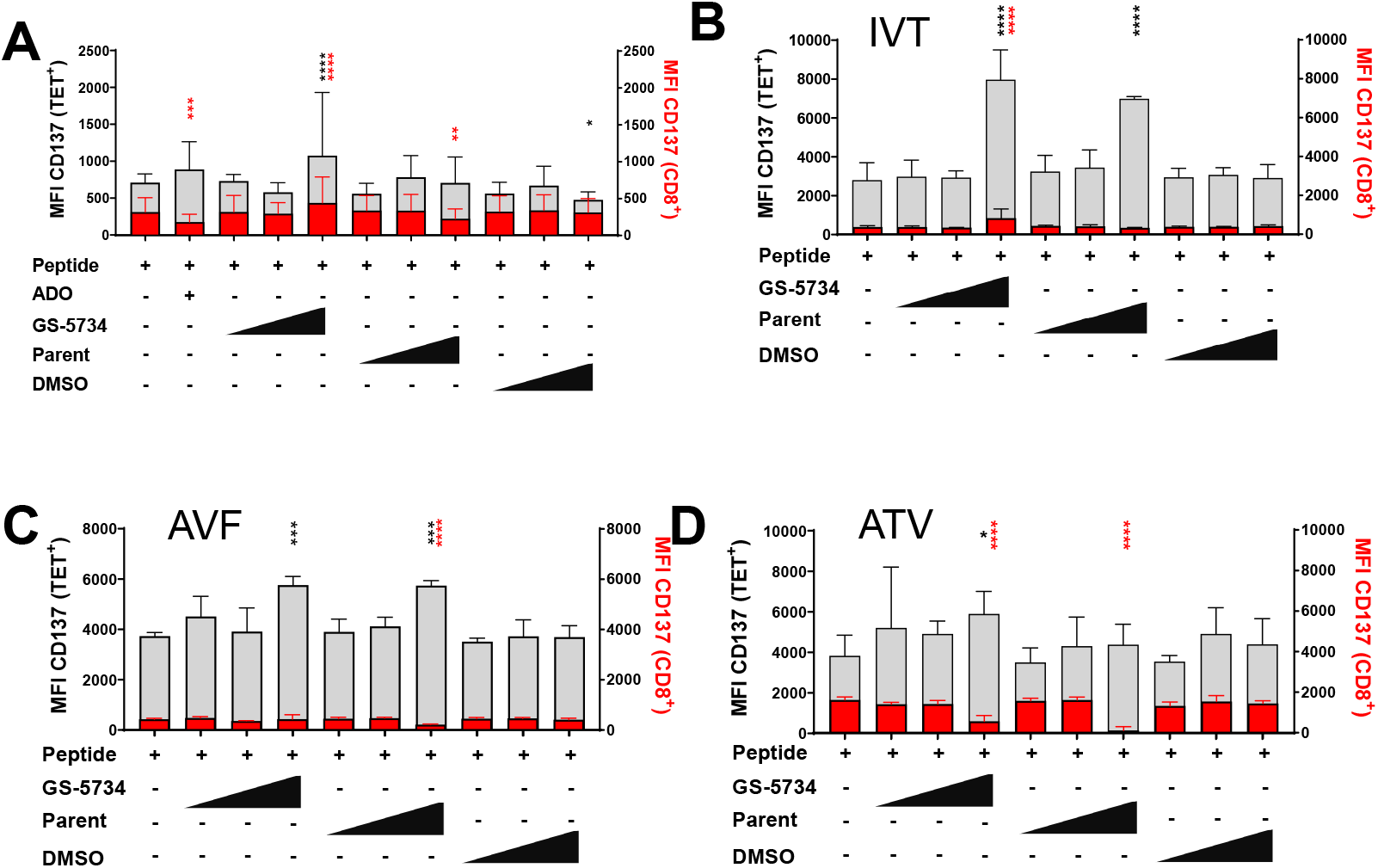
Effect of drug on T cell activation. A) HLA-A*02:01 Donor #1 PBMCs and B-D) HLA-A*11:01 Donor #3 PBMCs were treated and assayed on day 7 as described in Figure 1. Grey bars represent CD137 MFI in CD8^+^ tetramer positive T cells, red bars represent CD137 MFI in CD8^+^ tetramer negative T cells (bystander). ADO was omitted from donor #3 data due to low number of tetramer positive T cells. Data are representative of 4 (A) or 3 (B-D) independent experiments. Dunnett’s multiple comparison test relative to peptide-only treatment. *, p ≤ 0.05; **, p ≤ 0.01; ***, p ≤ 0.001; ****, p ≤ 0.0001.

## DISCUSSION

We have described a simple drug screening platform that can be utilized for preliminary assessment of potential off-target T cell effects occurring from small molecule treatment, as well as other therapeutic drug classes. The assay was originally designed to mimic one of the major aspects of the TME and thereby assess activity of candidate drug(s) to reverse adenosine-mediated suppression. The assay platform offers advantages in that it utilizes donor PBMCs representing a heterogeneous mix of most immune cell types. It also offers the advantage of a simple physiological T cell stimulation and the opportunity to assay effects on bystander cells. Here we show that the effects of high dose treatment with GS-5734 and GS-441524 on antigen-specific T cell expansion are consistent across donors and peptide stimulations with varied effects on antigen and non-antigen-specific T cell activation. Even though these two molecules are closely related in structure, their effect at high doses on antigen-specific T cell recall was very different. This illustrates the utility of this assay to screen drugs of varying structures as to whether memory T cell proliferation and function is unencumbered with drug administration. In theory, this assay is amenable to a variety of other drug classes such as those used in cancer immunotherapy ^18^ to monitor for immune-related adverse events (irAEs).

Here, we demonstrate the utility of this assay to investigate off-target effects on CD8^+^ memory T cells. However, this assay may also be used for assessment of off-target effects on CD4^+^ memory T cells. We predict enhanced utility of this assay by matching the relevant immunodominant recall peptide(s) with the appropriate investigational drug. In the case of SARS-CoV-2, this would entail securing PBMCs from donors who have had previous “experience” with SARS-CoV-2, stimulating them with relevant immunodominant peptide(s) and co-treating with drug candidate to assess off-target T cell effects. Furthermore, activation read-outs requiring earlier time points can be accomplished with donor PBMCs harboring detectable levels of tetramer positive T cells prior to expansion or by combining previously expanded cells with unexpanded as a source of fresh APCs.

## ACKNOWLEDGEMENTS

We thank Dr. Emma Chen (Crown Biosciences), Dr. Adel Benlahrech (ImmunoCore) and Dr. Reece Marillier (iTeos Therapeutics) for critical review of the manuscript.

## FOOTNOTE PAGE

Authors report no conflict of interest.

All research was funded by JSR Life Sciences LLC.

Corresponding author, Eden Kleiman, is currently located at Crown Bioscience Inc. (JSR sister company) at 16550 W Bernardo Dr Ste 525, San Diego, CA 92127. All lab work related to this submission was performed at JSR Life Sciences LLC at 1280 North Mathilda Avenue, Sunnyvale, CA 94089.

Current email addresses.

## Notes

### Competing Interest Statement

The authors have declared no competing interest.

### Summary of Updates

This version has been revised to accurately define GS-5734 (Remdesivir) as a prodrug of a nucleoside analog and not a nucleoside analog itself.

