## Supplemental Figure Legends for "Antigen-Specific T Cell Recall Assay To Screen Drugs For Off-Target Effects"

**Supplementary Figure 1**. Adenosine reduces cell proliferation. HLA-A*02:01 Donor #1 PBMCs were first incubated with proliferation dye and then treated with anti-CD3/anti-CD28 beads with or without ADO for 4 days. Three independent experiments are shown. Data were analyzed using Flowjo software proliferation modeling on live T cells. Displayed inside each histogram plot window is division index, percent divided, expansion index and number of cells per division.

**Supplementary Figure 2**. Adenosine does not decrease $Ki67^+$ MFI or the percentage of $Ki67^+$ cells from the tetramer T cell fraction. HLA-A*02:01 Donor PBMCs from Donor #1 were treated with CMV pp65 NLVPMVATV ± ADO and assayed for tetramer positive cell Ki67 staining on Day 7 using the TrueNuclear Transcription Factor Staining kit. A) Flow plots depict representative data showing unstimulated (U), Peptide stimulated (P) and P + ADO. Bar graphs depict the B) percentage of tetramer positive T cells that are Ki-67 positive and the C) MFI of tetramer positive T cell Ki67 displayed as a ratio relative to peptide treated cells. Results are from 6 independent experiments. Unpaired two-tailed t-test. ***, p = 0.0007. One experiment from another HLA-A*02:01 donor (#2) yielded similar results (not shown).

**Supplementary Figure 3**. ADO inhibits antigen-specific T cell recall and can be rescued by adenosine deaminase enzyme supplementation, and ADO suppression is not recapitulated by administration of adenosine breakdown metabolite inosine. A) HLA-A*02:01 Donor #1 PBMCs treated either with peptide alone (CMV pp65 NLVPMVATV), peptide + ADO, peptide + 10µg/mL adenosine deaminase (ADA) or a combination of peptide + ADO + ADA. Results from four separate experiments are expressed as the fold-change (FC) of tetramer positive T cells relative to peptide only treated wells. B) Recall experiment setup as in A, comparing addition of ADO versus ADO metabolite inosine (INO), both at the same concentrations. Results are from two independent experiments. Dunnett's multiple comparison test relative to peptide-only treated well. ****, $p < 0.0001$, *, $p=0.04$.

**Supplementary Figure 4**. Adenosine-mediated alterations in co-inhibitory receptor expression. HLA-A*02:01 Donor #1 PBMCs treated with 2.5µg/mL CEF peptide pool (ProMix CEF Peptide Pool, Proimmune Ldt) containing CMV pp65 NLVPMVATV for two experiments or 10µg/mL pure CMV pp65 NLVPMVATV peptide for two experiments. Cells were treated with either peptide/peptide cocktail (P) alone or with ADO. In these early experiments, ADO was administered on days 0, 2, 4 and 6. Surface expression of co-inhibitory receptors A) PD-1, B) LAG-3 and C) TIM-3 were assessed on CMV pp65 NLVPMVATV-specific $CD8^+$ T cells or tetramer negative $CD8^+$ T cells on day 7. Results are from 4 independent experiments. Two-tailed unpaired t-test analysis. *, $p < 0.05$.

**Supplementary Figure 5**. Adenosine stimulation increased the percentage of $CD14^+$ $CD11b^-$ cells and their surface expression of PD-1 co-inhibitory receptor ligands PD-L1/PD-L2 in Donor #1. HLA-A*02:01 Donor #1 PBMCs treated with or without 1mM ADO once for either 24 hours or 48 hours. Cells were assayed for (A-B) the percentage of $CD14^+$, $CD11b^-$ cells and (C-D) the surface expression of PD-L1/PD-L2 (same channel) on $CD14^+$, $CD11b^-$ cells. For C-D, grey bars depict the PD-L1/PD-L2 expression levels on $CD14^+$ $CD11b^-$ cells while black bars depict PD-L1/PD-L2 expression on T cells for comparison. Results are from 2 independent experiments for each time point. Two-tailed unpaired t-test analysis. **, $p \leq 0.01$; ***, $p \leq 0.001$; ****, $p \leq 0.0001$. Results from $CD14^+$ $CD11b^+$ cells did not show a significant difference in MFI (data not shown).

**Supplementary Figure 6**. Adenosine stimulation does not reduce IFNγ production or activation-induced TOX expression on previously expanded cells re-stimulated with peptide. HLA-A*02:01 Donor #1 PBMCs that were previously expanded (with NLVPMVATV peptide at least 7 days) were combined with unexpanded PBMCs of the same donor at a ratio of 2:1 on day 0. Cell co-cultures were either not re-stimulated or re-stimulated with 10µg/mL NLVPMVATV peptide (P) for 48 hours. Additionally, 1mM ADO was either added or not as indicated at the same time as peptide re-stimulation. A) representative plots showing IFNγ and TOX expression among tetramer positive T cells, except for black dots which are from CD3- cells. IFNγ plots with y axis tetramer and CD3 are meant to illustrate activation due to internalization of TCR (tetramer staining) and CD3. B-E) Bar graphs depicts IFNγ and TOX MFI and percentage of positivity within the tetramer T cell gate either stimulated with peptide (grey bars) or non-peptide-stimulated (black bars). Because tetramer positive cells from Donor #1 are only detectable by flow after day 5 post-peptide stimulation, this data is representative of the IFNγ and TOX expression that was induced 48 hours post-peptide stimulation on tetramer positive cells already present (expanded) as opposed to unexpanded PBMCs which will not have detectable tetramer positive cells at this time point. Staining was performed using TrueNuclear Transcription Factor kit with Brefeldin A treatment 4 hours prior to staining. Results are from 3 independent experiments. Data depicted as bar graph but two-tailed paired t-test analysis was performed using paired data points from each individual experiment. Statistical analysis only shown for peptide treated samples (grey bars), black bars displayed for reference. *, p = 0.01.

**Supplementary Figure 7**. Antigen-specific T cell recall is altered at high doses of GS-5734 and parent drug. HLA-A*02:01 Donor #1 PBMCs treated with CMV pp65 NLVPMVATV and assayed for tetramer positive cells on Day 7. ADO dissolved in complete media is denoted as $ADO^{aq}$. ADO dissolved in DMSO ($ADO^{DMSO}$), GS-5734 (DMSO) and parent drug GS-445124 (DMSO) were all given at ascending doses of 500nM, 5µM or 50µM. DMSO vehicle control was administered at 0.1%, 0.01% or 0.001% final. GS-5734 at 50µM is depicted as a grey striped bar to denote the toxicity observed. NECA and CGS-21680 (DMSO) were both given at ascending doses of 100nM, 1µM or 10µM. All treatments are normalized to peptide alone treatment (comparison using horizontal line at 1 on y axis). Data is from 4 independent experiments. The percentage of T cells that were antigen-specific ranged from 7-18%. Dunnett's multiple comparison test. ****, adjusted p value < 0.0001; ***, adjusted p value = 0.003; *, adjusted p value = 0.03.

**Supplementary Figure 8.** Cellular toxicity of drug treatments. HLA-A*02:01 Donor #1 PBMCs treated overnight with indicated drugs or DMSO vehicle. No peptide stimulation was used for this experiment. Indicated cell fraction (y axis label) was analyzed by Annexin V (AnV) vs 7AAD co-staining. Displayed are the single positive AnV cells (grey bars – early apoptosis) as well as the AnV positive, 7AAD positive cells (black bars – late stage apoptosis). Data is from 3 independent experiments. Data from all treatments are normalized to unstimulated. Dunnett's multiple comparison test. ***, adjusted p value < 0.0005; **, adjusted p value < 0.005; *, adjusted p value < 0.05.

**Supplementary Figure 9**. Cell viability is not significantly altered with drugs treatments, except for 50µM GS-5734. HLA-A*02:01 Donor #1 PBMCs treated with CMV pp65 NLVPMVATV along with various other treatments were assessed on day 7. A) FACS plots illustrating the scatter of cells from day 7 cultured cells. 50µM GS-5734 results in significant depletion of the lymphocyte gate. B) The percentage of $7AAD^+$ cells (among total cells) was assessed from Day 7 cultured cells. Data is from 5 independent experiments. Within each experiment, the percentage of $7AAD^+$ cells is normalized to peptide treated alone. Dunnett's multiple comparison test. *, p=0.02. The amount of cell death for 50µM GS-5734 treated cells, as assayed by 7AAD alone, is likely an underestimate as a significant proportion of the presumed non-viable small cells/debris induced by this treatment do not take up 7AAD dye but do stain for Annexin V (data not shown).

**Suppelmentary Figure 10**. Cell viability is not significantly altered with drug treatments. HLA-A*11:01 Donor #3 PBMCs treated with 10µg/mL of either A) EBV EBNA-3B peptide IVTDFSVIK, B) EBV EBNA4 peptide AVFDRKSDAK, or C) CMV pp65 peptide ATVQGQNLK along with various other treatments as in Supplementary Figure 7. The percentage of 7AAD positive cells (among total cells) was assessed on Day 7. Data is from 3 independent experiments. Within each experiment, percentage of $7AAD^+$ cells is normalized to peptide treated alone. Dunnett's multiple comparison test. As described in the figure before, the percentage of 7AAD$^{+}$ for 50µM GS-5734 is underestimated because small cells/debris induced by drug do not take up this dye.

**Supplementary Figure 11**. CD137 co-stimulatory surface marker used as a surrogate for activation status on day 7 of recall. Displayed is data from HLA-A*02:01 Donor #1 PBMCs treated with peptide only (NLVPMVATV) as a representative. CD137 expression non-overlaid histograms (bottom left) for CD3 negative cells (orange), CD3$^{+}$ CD8$^{+}$ tetramer negative cells (red) or CD3$^{+}$ CD8$^{+}$ tetramer positive cells (blue). Overlay plots include CD3, tetramer, 7AAD viability dye, CD4, CD8 and CD45RO.

**Supplementary Figure 12**. Effect of drug on T cell activation. HLA-A*02:01 Donor #1 PBMCs were treated as in Figure 1C and assayed for CD137 on day 7 as described in Figure 2. Grey bars represent CD137 MFI in tetramer positive T cells. Data are representative of 4 independent experiments. Dunnett's multiple comparison test relative to peptide-only treatment. *, $p \leq 0.05$; ****, $p \leq 0.0001$.
